# The Histopathology of *Cephenemyia stimulator*-Induced Nasopharyngeal Myiasis in Roe-Deers (*Capreolus capreolus*)

**DOI:** 10.1101/2023.11.23.568488

**Authors:** Irene Ortiz-Leal, Mateo V. Torres, Ana López-Beceiro, Pablo Sanchez-Quinteiro, Luis Fidalgo

## Abstract

Nasopharyngeal myiasis in European roe deer *(Capreolus capreolus)* is a pathological condition caused by the larval stages of *Cephenemyia stimulator*, a fly from the Oestridae family. These larvae reside in the host’s upper respiratory tract for months, inducing significant tissue damage and clinical symptoms. The lifecycle of *Cephenemyia stimulator* is complex, involving three larval stages before maturation into adult flies, with each stage contributing to the progressive pathology observed in the host. Despite their prevalence, the histopathological effects of these larvae in the nasopharyngeal and nasal cavities have been understudied. Our study fills this knowledge gap by providing a detailed histopathological analysis of the affected tissues, using various staining techniques to reveal the extent and nature of the damage caused by these parasitic larvae. This histopathological examination reveals significant alterations within the nasopharyngeal mucosa and nasal cavity, including erythematous changes, mucosal metaplasia, fibrosis, and tissue necrosis. Parasitic cysts and eosinophilic infiltration further characterize the infestation’s impact, compromising not only the mucosal integrity but also potentially the olfactory function of the affected animals. This research is crucial for understanding the impact of myiasis on the health of roe deer populations and could have significant implications for wildlife management and conservation.

## INTRODUCTION

Nasopharyngeal myiasis in ruminants is caused by three genera of flies from the Oestridae family: *Pharyngomya, Oestrus*, and *Cephenemyia*. Of the 8 species of the genus *Cephenemyia* that cause myiasis, *Cephenemyia stimulator* is particularly notable in roe deers. The adults are short-lived flying insects, whereas the larvae spend several months in the upper respiratory tract, responsible for the lesions and symptoms observed.

The *Cephenemyia stimulator* flies undergo complete metamorphosis through three larval stages (L1, L2, L3), each with size increase and morphological changes. The L3 larva leaves the host and falls to the ground to become a pupa, a phase characterized by a chitinous, thick, and dark skin that allows it to survive buried. The adult emerges from the pupa and has a brief lifespan dedicated to reproduction, although this period can be extended under adverse climatic conditions through diapause. Sugár (1974) described how the first-stage larvae found in the nasal mucosa are immobile in the ethmoid turbinates. The second-stage larvae were found in the nasal turbinates as well as in the pharyngeal recesses, while the third-stage larvae were located in the pharyngeal recesses, in the nasal cavities, and in the larynx.

The pharyngeal recesses are greatly enlarged—five to six times or even more than the normal size—although the specific mechanism causing this distension is unknown (Sugár, 1974; Cogley, 1987). After the death of the animal, as Sugár (1978) described, some L1 larvae may move towards the throat and other deeper locations of the respiratory system, in the same way that the L3 larvae leave their location to move outward through the nostrils or towards the trachea or esophagus.

The larvae feed primarily on mucus and respiratory secretions, obtained thanks to the irritating effect of the numerous spicules present in their cuticle. The mouth hooks, which allow them to cling to the mucosa, further injure and irritate it. They feed through a mechanism of extracorporeal digestion, in which they release their digestive enzymes into the environment and also excrete the waste products of their metabolism, causing irritation to an already injured mucosa (Sugár, 1974; Cogley, 1987). As a result of the foregoing, the presence of the larvae causes catarrhal inflammation of the nasopharyngeal mucosa, although macroscopically ulcerative or bleeding lesions are not observed (Tabouret et al., 2001, 2003; Angulo-Valadez et al., 2010). The infestation results in a clinical picture of behavioral alterations, deterioration of body condition and antler development, respiratory symptoms, loss of the ability to flee from predators, and a higher predisposition to other concomitant diseases.

This specific parasitosis in European roe deer was first described in 1970 in Poland, and it quickly spread to other regions; currently, it is known to affect at least 12 countries (Kusak et al., 2012; Žele Vengušt et al., 2021; Flis et al., 2021). The distribution of *Cephenemyia stimulator* is widespread and exclusive across Europe, notably overlapping with the range of the roe deer. In Spain, this disease was first described in 2001 (Notario and Castresana, 2001). However, the earliest complete data on its incidence and distribution in Spain are from 2013 (Fidalgo et al., 2013). Research on the prevalence of this parasitism in roe deer is scarce and often refers to a limited number of specimens, affecting the quality of the results. A prominent study in Hungary, with a large number of deer examined (956), showed an incidence of 34.6%, indicating that younger animals are most vulnerable to infection, followed by adult males and finally adult females (Király and Egri, 2007).

Despite the time that infestations have been impacting European roe deer populations, to our knowledge, there is not any histopathological study that captures the effects of the lesions in both the pharyngeal and nasal cavities. In an effort to fill this gap, we have carried out a histopathological examination of a substantial number of samples, which confirms the extent of macroscopic damage in the both the olfactory and pharyngeal mucosae, as well as identifies the primary histopathological alterations induced by the myiasis using various histological staining techniques.

## MATERIALS AND METHODS

### Samples

For the study, 6 roe deer with larvae of *Cephenemyia stimulator* in various stages of development were selected, along with 2 healthy deer, free from the disease, collected during the years 2021 and 2022. The deer originated from the official wildlife recovery centers of Galicia (northwest Spain) and from legally authorized hunting activities. Immediately after the death of the animal, the head is collected and transported refrigerated to be processed within 8 hours post-mortem.

### Pathological study

A thorough postmortem investigation was performed on all the samples received. Representative tissues from the nasal cavity and the pharynx mucosa were preserved in Bouin’fluid for 24 hours before transferring them to 70% ethanol.

Fixed tissues were dehydrated through baths in graded alcohol and xylene, followed by embedding in paraffin; 6-μm-thick sections of the paraffin-embedded tissues were rehydrated and stained with hematoxylin and eosin, PAS and Alcian-blue stains. The detailed procedures are described in Barrios et al. (2014) and Ortiz-Leal et al. (2020).

Digital photographs were taken using a digital camera from Karl Zeiss, model MRc5 Axiocam, connected to a microscope of the same brand, model Axiophot. The Adobe Photoshop CS4 software (Adobe Systems, San Jose, CA) was used for those images that required digital enhancement of brightness, contrast, and white balance. However, no enhancements, additions, or relocations of the image features were made.

## Data availability

The underlying data will be made available upon request.

## RESULTS

The nasal and nasopharyngeal caivties were the primary focus of the histological investigation of the *Cephenemyia* infestation since it was there that the parasite’s larvae in various developmental stages were found. The nasal turbinates of the nasal cavity and, more severely, the mucosa lining the pharyngeal pouch of the nasopharynx were the particular sites that were injured.

## Macroscopic study (Fig. 1)

The localization of *Cephenemyia* larvae is concentrated in the recess formed by the pharyngeal pouch or pharyngeal recess (also known as Rosenmüller’s fossa in human medicine). This consists of a tiny epithelial pouch located in the caudodorsal part of the medial mucosa of the nasopharynx.

**Figure 1.**
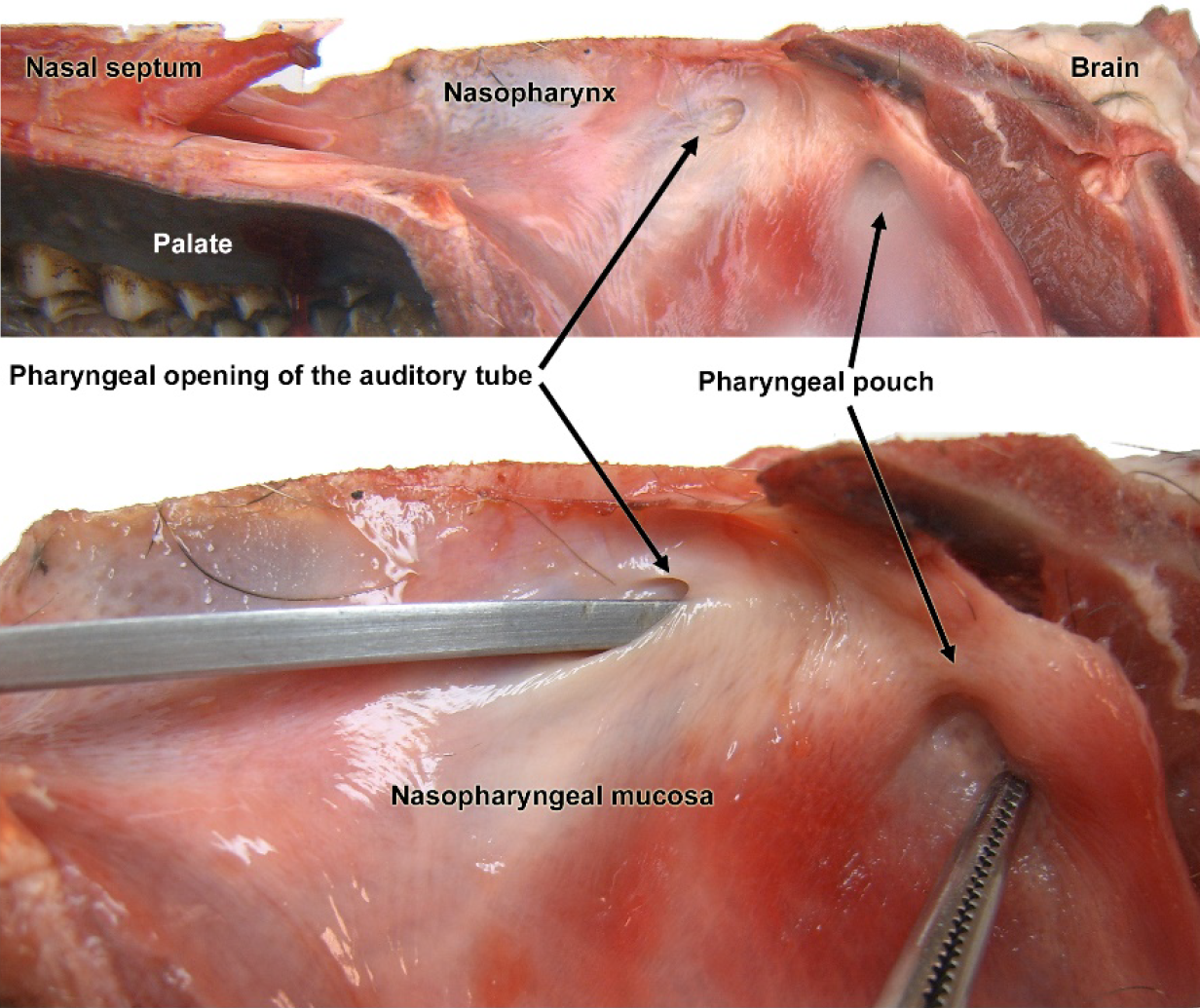
N**asopharynx mucosa of the roe deer**. Dissection of the medial mucosa of the nasopharynx showing the topographic relationships of the pharyngeal recess and the pharyngeal opening of the auditory tube.

It has a direct topographical relationship with the pharyngeal opening of the auditory tube: the communication between the middle ear and the nasopharynx. The appearance and color of the mucosa is reddish, with moderate mucosity, stemming from the secretory activity of the mucous glands present in its own lamina.

During parasitic infestation, the larvae of *Cephenemyia* occupy the cavity of the pouch, causing a notable distension of the mucosa, which is accompanied by a thickening of the wall, the formation of edema, and an extensive secretion of mucus (Fig. 2).

**Figure 2.**
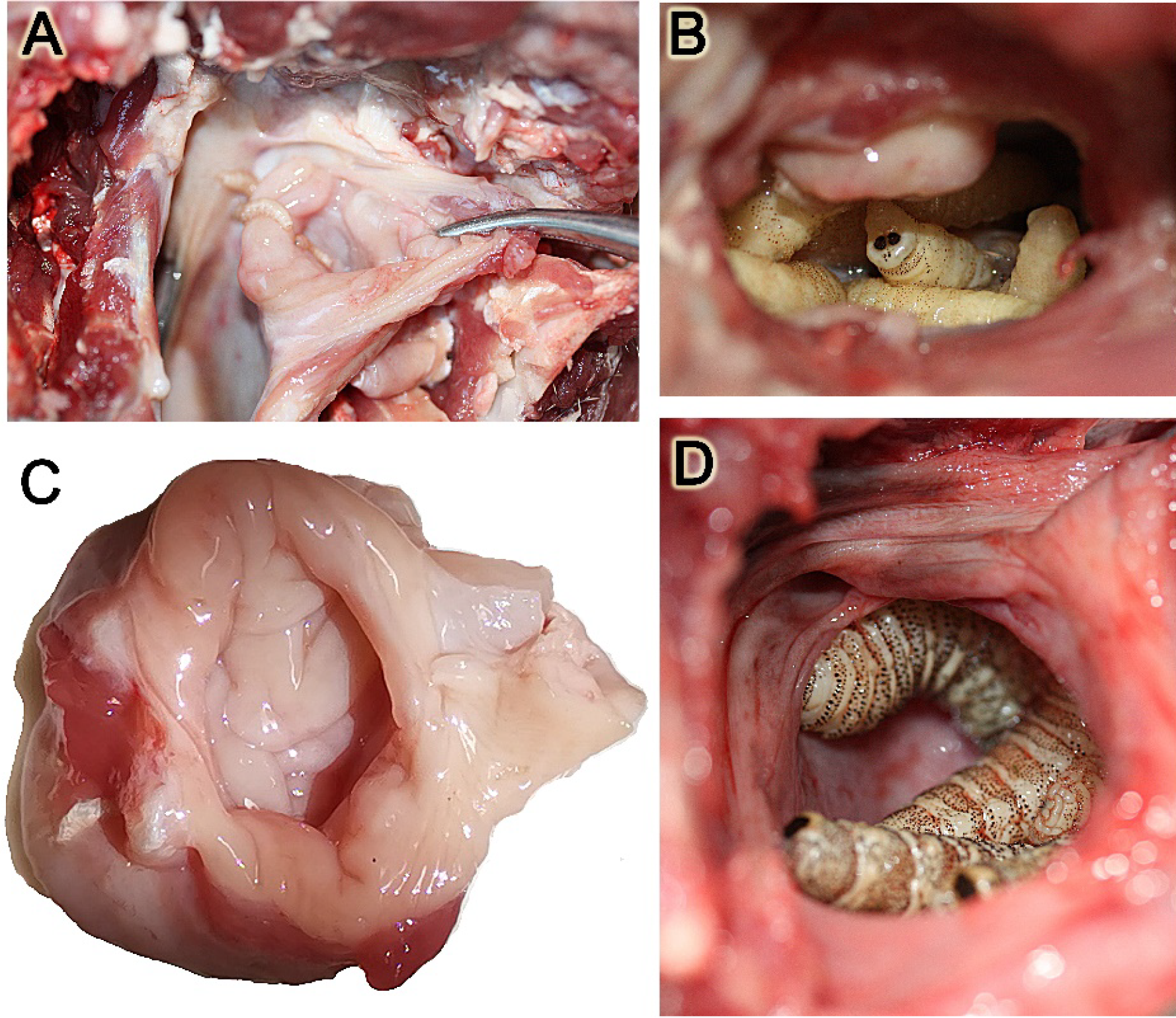
Macroscopic images of the pharyngeal pouch in situations of maximum infestation by Cephenemyia larvae. The inflammatory reaction is superficially reflected in a thickening of the mucous lining and the production of an abundant mucous secretion.’

### Histopahtology

In healthy animals, the nasopharyngeal mucosa has an intense reddish-brown coloration and a smooth, slippery surface covered with a transparent mucus (Fig. 3A). Microscopically (Fig. 3B), the mucosa is characterized by the integrity of its strata and the normal appearance of the epithelial cells that form the mucosal lining (Fig. 3C). Likewise, the lamina propria shows a remarkable development of collagen fibers, smooth muscle fibers and extensive glandular accumulations that are characterized by being both PAS-positive (Fig. 3C, D) and Alcian Blue-positive (Fig. 3E).

**Figure 3.**
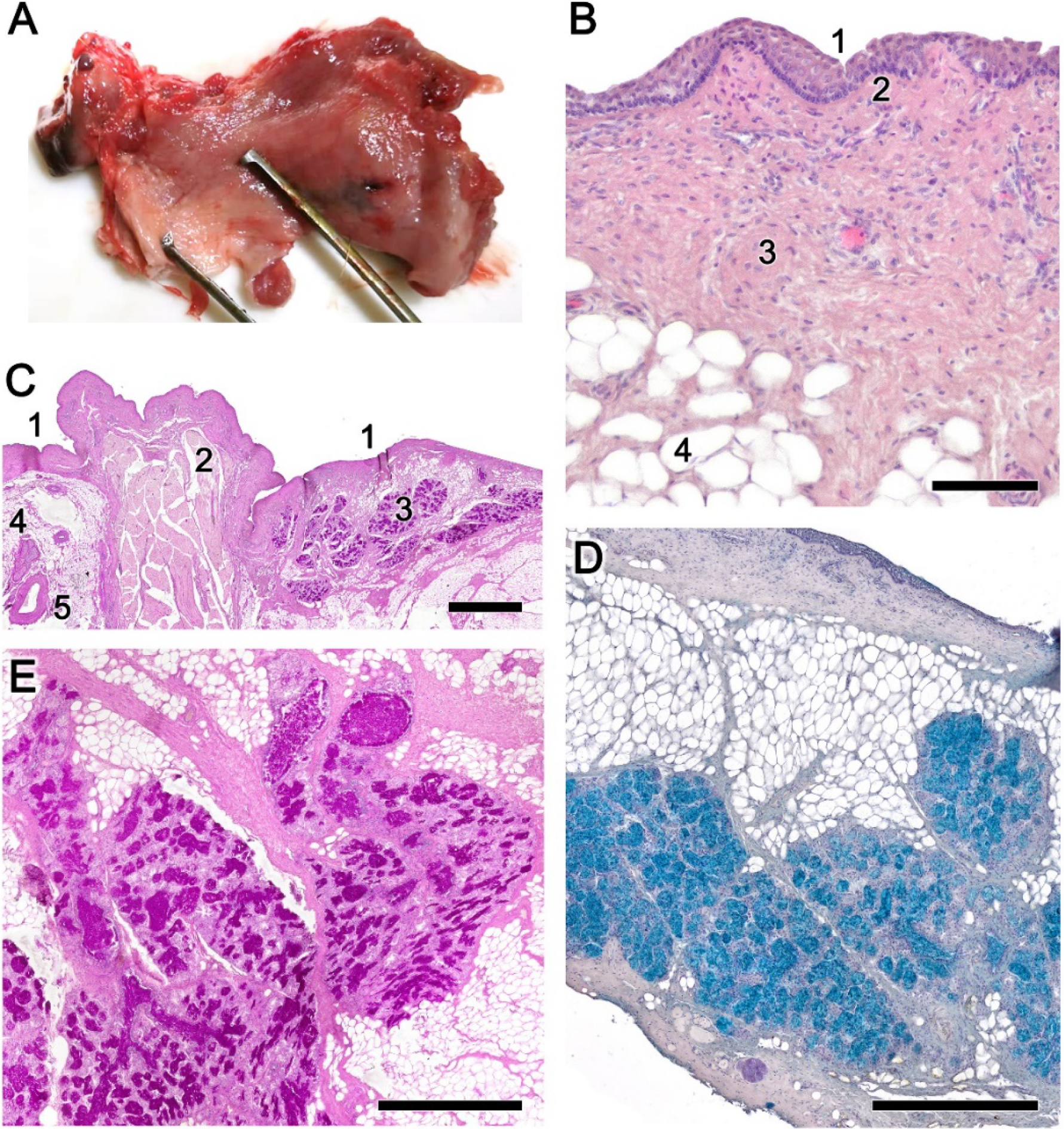
Normal anatomy and histology of the nasopharynx mucosa of the roe deer. **A**. Macroscopic anatomy of the nasopharyngeal mucosa showing the normal intense reddish-brown coloration and the smooth and uniform surface, devoid of folds. **B**. Microscopically, the mucosa is characterized by the integrity of its strata and the normal appearance of the epithelial cells that form the mucosal lining. 1, mucosal epithelium; 2, Basal stratum; 3, Collagen fibers; 4. Adipose tissue. **C**. The thickness of the epithelial covering (1) is variable depending on the area considered. The lamina propria shows a remarkable development of smooth muscle fibers (2), glandular accumulations (3), nerves (4) and arteries (5). **D and E**. Glandular acini PAS and Alcian Blue-positive, **r**espectively. Staining: Hematoxylin-eosin: B; PAS, C, E; Alcian blue: D. Scale bars: C: 1 mm; D, E: 500 µm; B: 100 µm.

In infested animals, the mucosa presents a more erythematous appearance, with hemorrhagic areas, abundant folds and pits and less mucus (Fig. 4A). Histologically (Fig. 4B, C), severe mucosal injury is observed, with a loss of epithelial and glandular cells and replacement of the lamina propria by a dense infiltrate of metaplastic cells, in a fibrous connective tissue matrix. The deeper layers show tissue necrosis.

**Figure 4.**
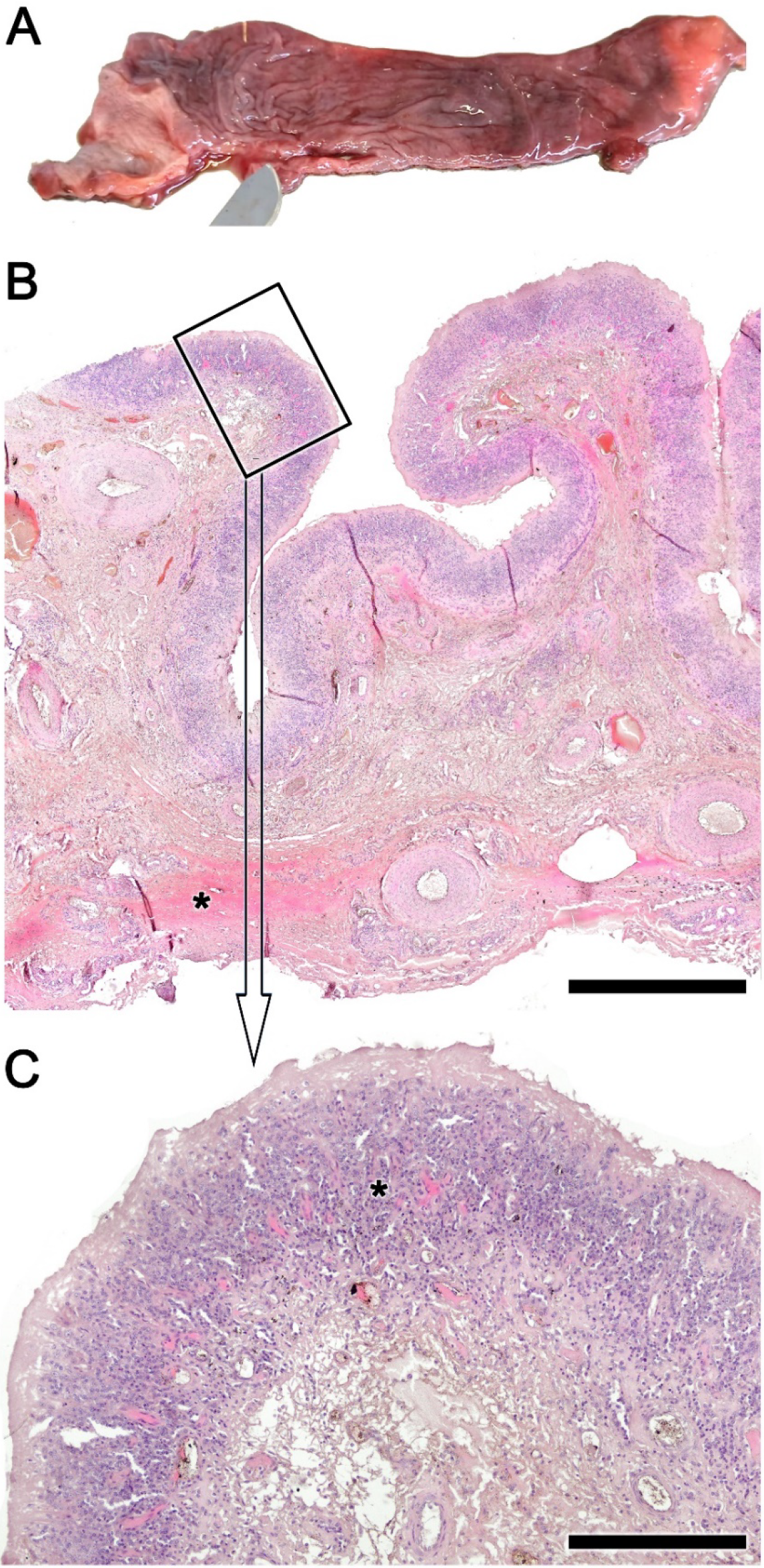
Anatomical and histopathological findings in the nasopharynx mucosa of the roe deer infested by *Cephenemyia*. **A**. The mucosa of infested animals presents an erythematous appearance, with hemorrhagic areas, abundant folds and pits and less mucus. **B**. Microscopically, a severe mucosal injury is observed, with a loss of epithelial and glandular cells and replacement. The deeper layers show tissue necrosis (asterisk). **C**. Higher magnification of the inset in B show in the lamina propria a dense infiltrate of metaplastic cells, in a fibrous connective tissue matrix (asterisk). Staining: Hematoxylin-eosin. Scale bars: B: 1 mm; C: 250 µm.

PAS staining allows for recognition of the high degree of mucosal fibrosis and identification of broad bands of necrosis in the submucosa (Fig. 5A). At higher magnification, it is observed how the fibrous tissue forms a trabecular network that encompasses metaplastic cells (Fig. 5B). In the arteries supplying the mucosa there is a prominent subintimal fibrosis with unfolding of the elastic fibers (Fig. 5C).

**Figure 5.**
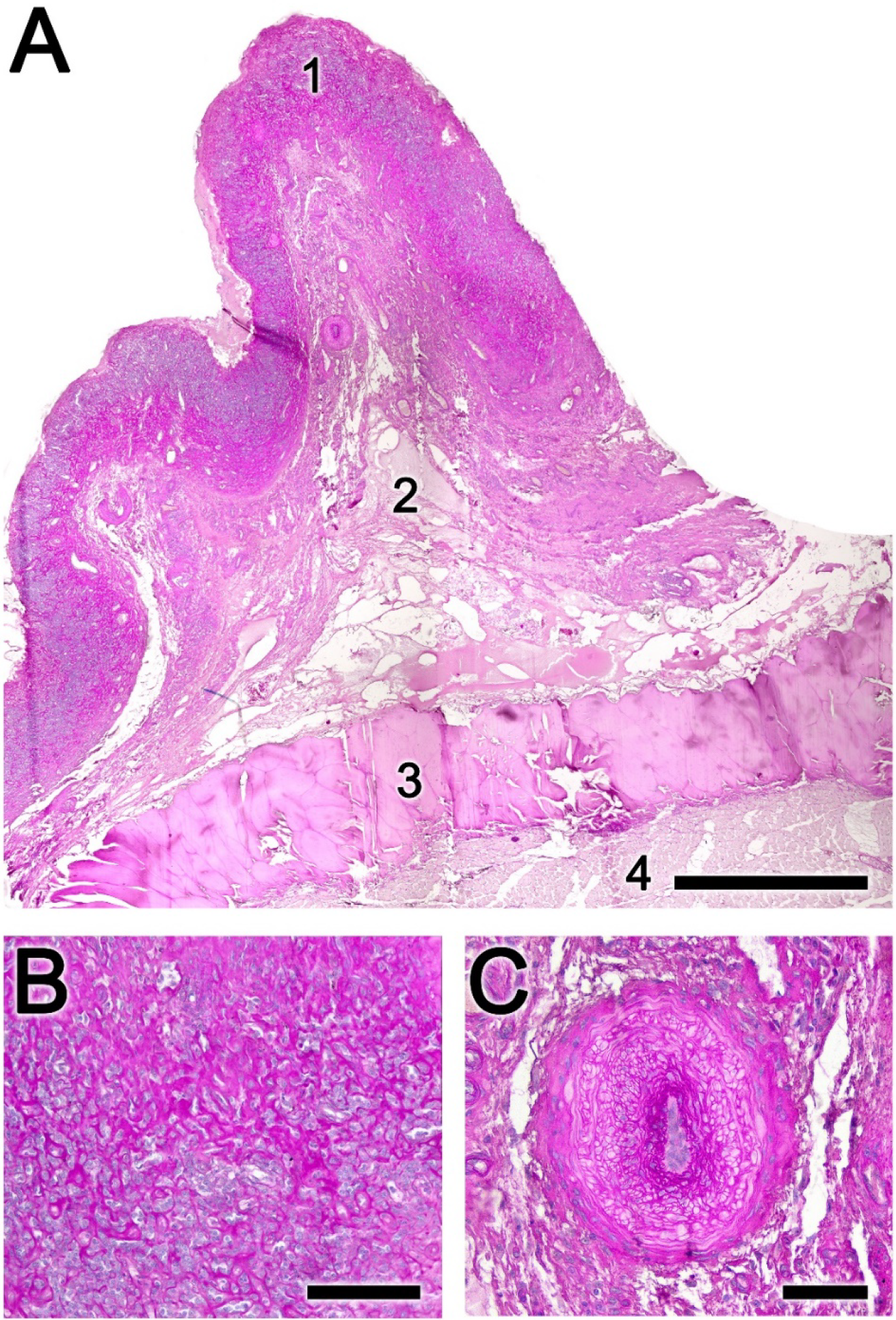
Histopathological findings in the nasopharynx mucosa of the roe deer infested by *Cephenemyia*. **A**. There is a high degree of mucosal fibrosis and broad bands of necrosis in the submucosa. 1, mucosal epithelium; 2, lamina propria; 3, submucosa; 4, muscular layer. **B**. The fibrous tissue forms a trabecular network that encompasses metaplastic cells. **C**. The arteries supplying the mucosa present a prominent subintimal fibrosis with unfolding of the elastic fibers. Staining: PAS. Scale bars: A: 1 mm; B: 250 µm; C: 100 µm.

Alcian blue staining reveals the undifferentiated nature of the metaplastic cells (Fig. 4A). Parasitic cysts in the muscle fibers, presumably *Sarcocystes* (Figs. 6B, C), and local foci of necrosis (Fig. 6D) are common findings.

**Figure 6.**
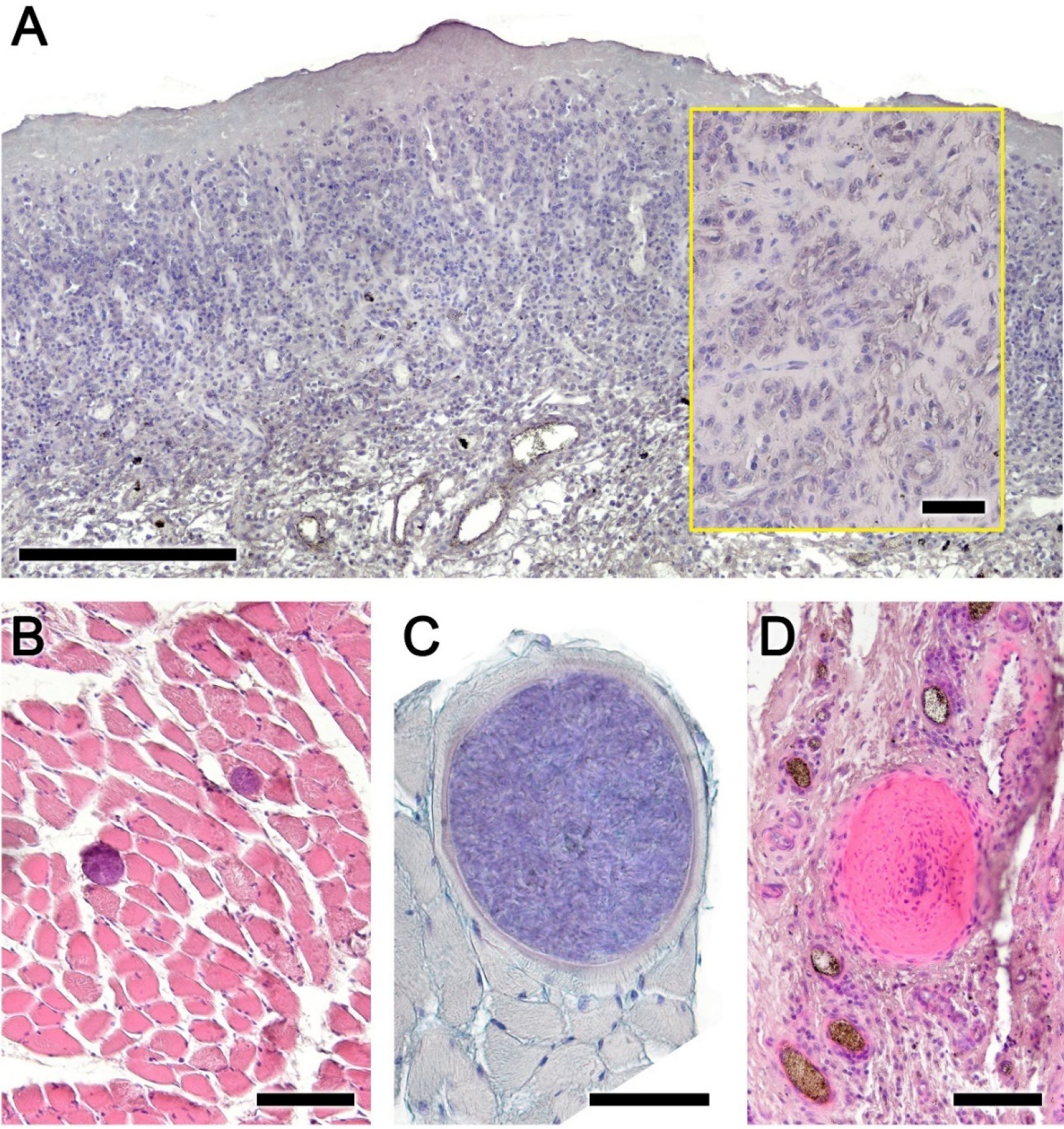
Histopathological findings in the nasopharynx mucosa of the roe deer infested by *Cephenemyia*. **A**. Alcian blue staining reveals the undifferentiated nature of the metaplastic cells. At higher magnification in the box. **B**. Parasitic cysts in the muscle fibers are usually observed. **C**. One cyst at higher magnification and stained with alcian blue. **D**. Local foci of necrosis are also common findings. Staining: A, C: Alcian blue; B, D: Hematoxylin-eosin. Scale bars: A: 250 µm; B, C, D: 100 µm; A inset: 50 µm.

It is common for a high degree of infestation (Fig. 7A) to cause extensive distension of the pharyngeal pouch, which accentuates the mucosal lesion resulting in intense mucosal metaplastic transformation (Fig. 7B). Degeneration and atrophy of muscle fibers, fibrosis in the lamina propria and intense metaplasia and fibrosis in the superficial zone are also observable (Fig. 7C, D).

**Figure 7.**
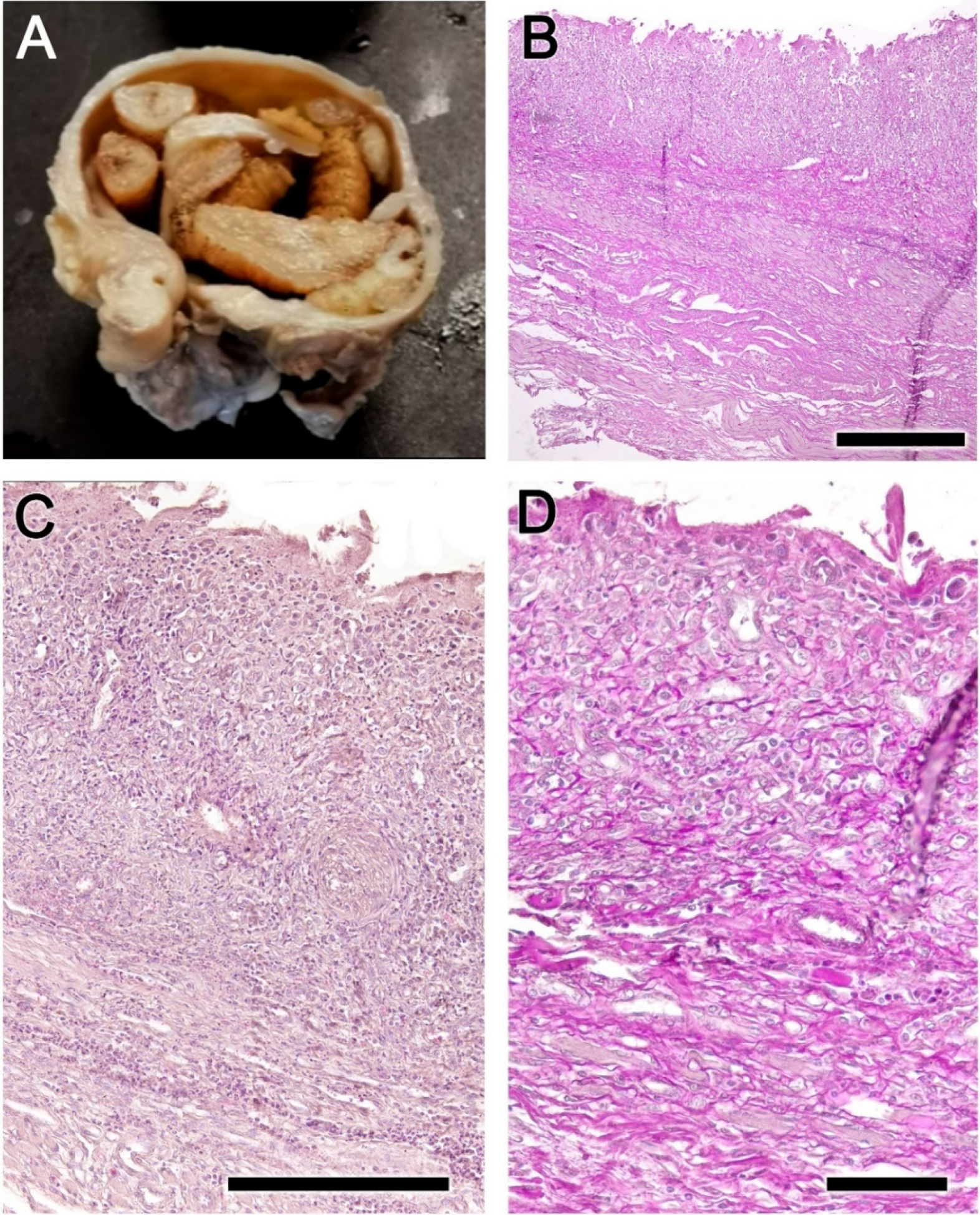
Histopathological findings in a high degree of infestation by *Cephenemyia* of the nasopharynx mucosa of the roe deer. **A**. The high quantity of larvae causes an extreme distension of the pharyngeal pouch. **B**. In these cases the metaplastic transformation in the mucosa is very intense. **C** and **D**. With both hematoxylin-eosin and PAS stainings, it is observed a degeneration and atrophy of muscle fibers, fibrosis in the lamina propria and intense metaplasia and fibrosis in the superficial. Staining: B, D: PAS; C: Hematoxylin-eosin. Scale bars: B: 500 µm; C: 250 µm; D: 100 µm.

### Nasal cavity

The lesions in the nasal mucosa of roe deer are also severe, compromising the animal’s olfactory function. The sampled area is shown in Figure 8. Both the olfactory mucosa (Figs. 9A-D) and the respiratory mucosa (Figs. 9E-F) show vacuolization and epithelial degeneration, with a loss of epithelial layering and a conspicuous eosinophilic infiltration. In the respiratory mucosa, significant vacuolization is also observed; however, the cellular profiles are not so much altered as in the olfactory mucosa.

**Figure 8.**
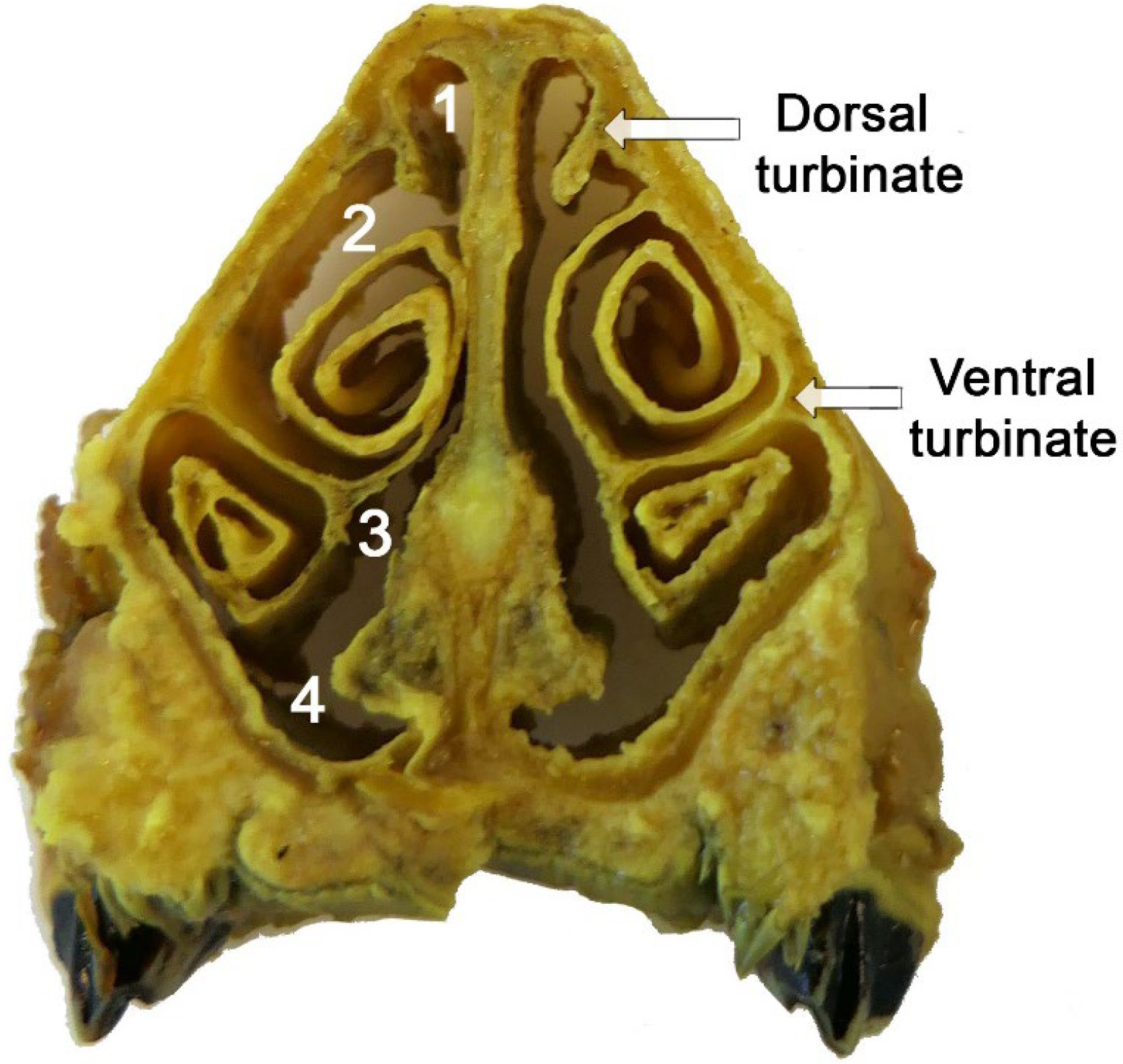
Transverse section of the nasal cavity of the roe deer. 1, dorsal meatus, 2, middle meatus, 3, common meatus, 4, ventral meatus.

**Figure 9.**
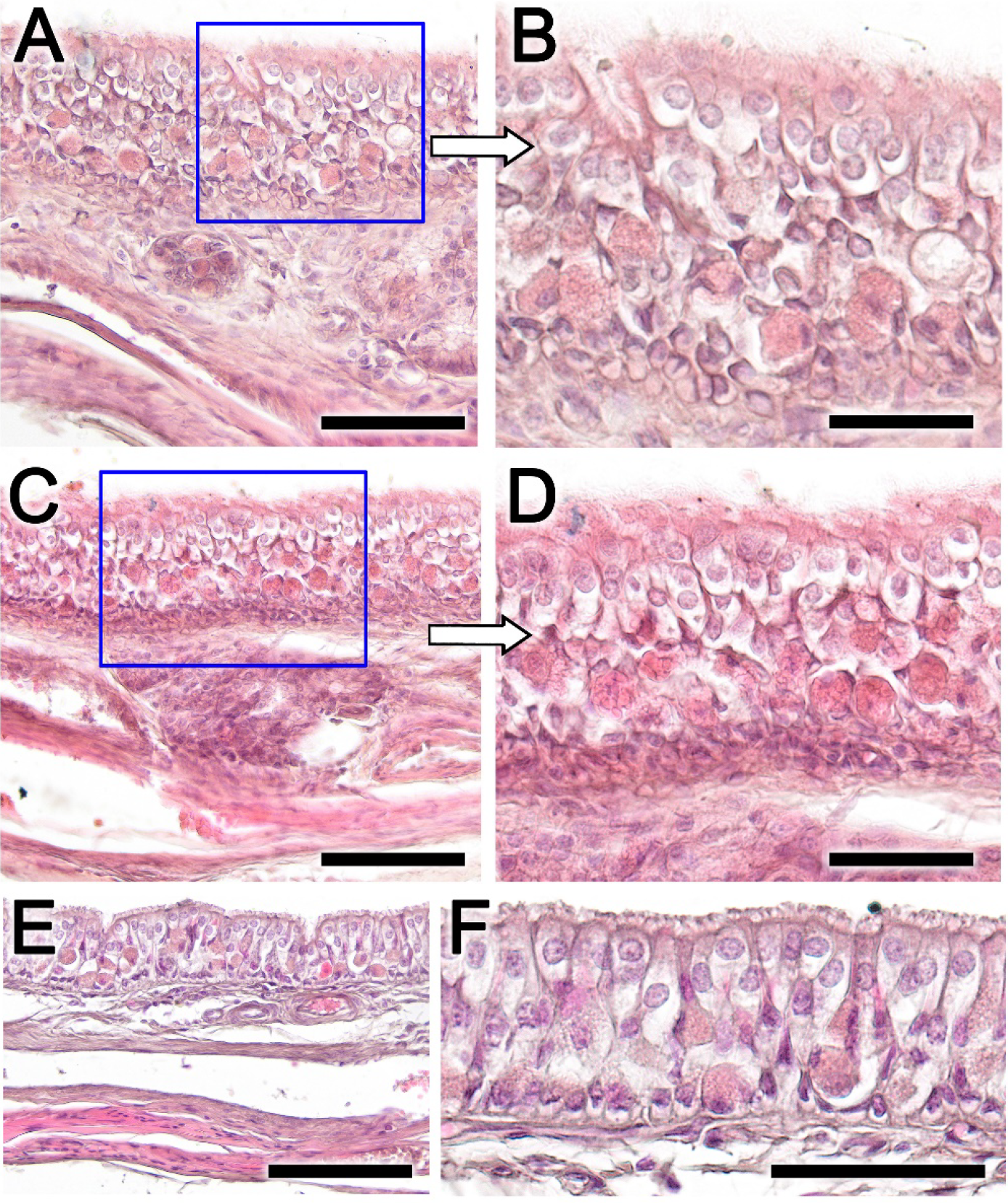
Histopathological findings in the olfactory and respiratory mucosa of the nasal cavity in the roe deer infested by *Cephenemyia*. **A** and **C**. General view of the olfactory mucosa in transverse sections at the level of the dorsal and ventral turbinates. The boxes, magnified **B** and **D** show vacuolization and epithelial degeneration, with a loss of epithelial layering and a conspicuous eosinophilic infiltration. **E** and **F**. In the respiratory mucosa, vacuolization is also observed; however, the cellular profiles are not so altered as in the olfactory mucosa. Staining: Hematoxylin-eosin. Scale bars: A, C, F: 100 µm; E, F: 50 µm.

## DISCUSSION

Currently, there is significant concern regarding the rapid spread and impact on wild roe deer populations of infestations by *Cephenemyia* larvae. This phenomenon, emerging and consolidating recently, has not been extensively studied, particularly in terms of its morphological and histopathological consequences. To our knowledge, beyond the work by Thomas Cogley (1987), who describes the pathologic changes induced by larvae of *Cepheneniyia* within the retropharyngea recesses of black-tailed deer (*Odocoileus hemionus columbianus*), there are no studies addressing the histopathology of the infestation. Such studies are critical to objectively assess the severity and implications of these infestations. Consequently, as we embark on this discussion, it’s important to note the limited body of work available for comparison and validation of the data we have gathered.

From a macroscopic point of view, the lesions we observed in the pharyngeal pouches of the roe deer are consistent with those described by Cogley (1987) in the black-tailed deer. Whereas under normal conditions the pharyngeal pouch corresponds to a small elliptical slit, infestation by Cephenemyia results in large circular dilations which contain dozens of larvae. In healthy specimens of roe deer, the mucosa exhibits robust anatomical features, such as well-defined epithelial layers and a well-vascularized lamina propria with dense collagenous and glandular structures. However, infestation precipitates drastic transformations. The normally reddish-brown, smooth nasopharyngeal mucosa becomes erythematous, hemorrhagic, and irregular in infested deer, accompanied by a notable decrease in mucus production.

Histologically, we have found in the roe deer how the nasopharyngeal cavity, specifically the nasal turbinates and the mucosa lining the pharyngeal pouch, endured pronounced histological changes due to *Cephenemyia* infestation. The degree of alteration we found in the pharyngeal mucosa of our specimens is much more severe than that observed by Cogley (1987) in the pharyngeal pouch the black tailed deer. Regarding the nasal turbinates, they were not included in Cogley’s study.

This morphological disruption caused by *Cephenemyia* infestation correlates with severe histological damage observed microscopically. A marked decrease in epithelial and glandular cell integrity, and a consequential replacement with fibrous connective tissue, underscores a pronounced inflammatory response. The application of PAS staining highlighted extensive mucosal fibrosis and submucosal necrosis, revealing a fibrous trabecular network engulfing metaplastic cells and suggesting a deep-seated tissue remodeling. Concurrently, arterial changes, including subintimal fibrosis, indicate a compromised blood supply, potentially exacerbating the local tissue distress. Alcian blue staining confirmed the presence of undifferentiated metaplastic cells, suggesting a loss of specific cellular function and further supporting the observed degenerative changes. Additionally, the discovery of parasitic cysts intermingled within the muscle fibers provides evidence of secondary parasitic involvement, potentially exacerbating the myiasis-induced lesions.

These lesions are more severe than those observed by Cogley in the black tail deer, who describes cellular desquamation with loss of the mucosal epithelium and inflammatory infiltration in the lamina propria, and glandular degeneration in the superficial area. In both cases, we agree on not finding purulent inflammation as well as the absence of neutrophils.

The nasal cavity lesions were equally severe, with evident vacuolization and degeneration of both the olfactory and respiratory mucosa, indicative of the larval invasion’s impact on sensory and respiratory functions. These findings corroborate the clinical observations of behavioral changes and compromised body condition in the infested deer, which could be partly due to disrupted olfactory cues affecting foraging and predator avoidance behaviors. These lesions in the olfactory mucosa have not, to our knowledge, been previously described in infestations with *Cephenemyia*.

Collectively, our results underscore the profound histopathological impact of *Cephenemyia stimulator* infestation on European roe deer. The extensive mucosal and submucosal alterations, including metaplastic and fibrotic changes, tissue necrosis, and secondary parasitic infections, delineate a severe condition that could have substantial implications for the health and survival of affected deer populations. These findings highlight the need for integrated pest management strategies and further research into the pathophysiological mechanisms underlying nasopharyngeal myiasis in wildlife.

## ACKNOWLEDGEMENTS

The authors wish to thank to the wildlife recovery centers from Galicia, and to ‘Dirección Xeral de Patrimonio Natural (Consellería de Medio Ambiente e Ordenación do Territorio, Xunta de Galicia)’, and ‘Federación Galega de Caza’ for having authorized and facilitated the sampling of the animals.

